# Reduced representation sequencing to understand the evolutionary history of Torrey pine (*Pinus torreyana* Parry) with implications for rare species conservation

**DOI:** 10.1101/2021.07.02.450939

**Authors:** Lionel N. Di Santo, Sean Hoban, Thomas L. Parchman, Jessica W. Wright, Jill A. Hamilton

## Abstract

Understanding the contribution of neutral and adaptive evolutionary processes to population differences is often necessary for better informed management and conservation of rare species. In this study, we focused on *Pinus torreyana* Parry (Torrey pine), one of the world’s rarest pines, endemic to one island and one mainland population in California. Small population size, low genetic diversity, and susceptibility to abiotic and biotic stresses suggest Torrey pine may benefit from inter-population genetic rescue to preserve the species’ evolutionary potential. We leveraged reduced representation sequencing to tease apart the respective contributions of stochastic and deterministic evolutionary processes to population differentiation. We applied these data to model spatial and temporal demographic changes in effective population sizes and genetic connectivity, to assess loci possibly under selection, and evaluate genetic rescue as a potential conservation strategy. Overall, we observed exceedingly low standing variation reflecting consistently low effective population sizes across time and limited genetic differentiation suggesting maintenance of gene flow following divergence. However, genome scans identified more than 2000 SNPs candidates for divergent selection. Combined with previous observations indicating population phenotypic differentiation, this indicates that natural selection has likely contributed to population genetic differences. Thus, while reduced genetic diversity, small effective population size, and genetic connectivity between populations suggest genetic rescue could mitigate the adverse effect of rarity, divergent selection between populations indicates that genetic mixing could disrupt adaptation. Further work evaluating the fitness consequences of inter-population admixture is necessary to empirically evaluate the trade-offs associated with genetic rescue in Torrey pine.

## Introduction

Conservation biology aims to preserve rare species and determine the appropriate management strategies necessary for long-term persistence and maintenance of evolutionary potential (Di Santo & Hamilton, 2020; Ralls et al., 2018; Swarts, Sinclair, Krauss, & Dixon, 2009; Young, Brown, & Zich, 1999). Rare species may have reduced effective population sizes (Ne), impeding populations’ ability to adapt to change (e.g., Ne < 1000) or increasing probability of inbreeding (e.g., Ne < 100) (Frankham, Bradshaw, & Brook, 2014), ultimately increasing the risk of local extirpation. Combined, rarity and isolation are often associated with stochastic loss of genetic variation (Aguilar, Quesada, Ashworth, Herrerias□Diego, & Lobo, 2008; Hague & Routman, 2016; Young, Boyle, & Brown, 1996). Genetic rescue is one conservation strategy that has been successfully used in both animals and plants to mitigate consequences of severely reduced genetic diversity (Bossuyt, 2007; Hedrick, Peterson, Vucetich, Adams, & Vucetich, 2014; W. E. Johnson et al., 2010; Madsen, Shine, Olsson, & Wittzell, 1999; Westemeier et al., 1998; Willi, Van Kleunen, Dietrich, & Fischer, 2007). Genetic rescue introduces or restores gene flow between populations to alleviate the fitness consequences of inbreeding through the introduction of genetic variation. However, while rare species may exhibit small effective population sizes and reduced adaptive potential, disruption of local adaptation may lead to outbreeding depression, or reduced fitness of progeny following admixture between genetically differentiated lineages (Hufford & Mazer, 2003). Thus, the contribution of natural selection to the evolution of population genetic differences is a consideration for genetic rescue, as it may ultimately lead to migrants or translocated individuals being maladapted (Lowry, Rockwood, & Willis, 2008; Nosil, Vines, & Funk, 2005). For rare species conservation, an understanding of contemporary effective population size is therefore required to assess immediate genetic threats to population persistence. Following these threats, however, understanding the history of population connectivity, the distribution of genetic variation within and among populations, and the role of selection in shaping population differences will be critical to informed management decisions.

In addition to guiding conservation management strategies, understanding rare species’ demographic and evolutionary history may prove valuable to optimizing strategies necessary to preserve neutral and nonneutral genetic diversity *ex situ*. *Ex situ* conservation collections, or the preservation of species outside their natural range of occurrence, can complement *in situ* conservation strategies (Cavender et al., 2015; Potter et al., 2017; Pritchard, Fa, Oldfield, & Harrop, 2012), providing a critical resource for the preservation of genetic variation, restoration or reintroduction (Guerrant Jr, Havens, & Vitt, 2014; Potter et al., 2017). *Ex situ* sampling designs traditionally rely on neutral population genetic structure to guide sampling decisions (Caujapé-Castells & Pedrola-Monfort, 2004; Gapare, Yanchuk, & Aitken, 2008; Hoban, 2019; Hoban & Schlarbaum, 2014). However, concerns exist regarding the sole use of neutral genetic variability for species conservation, as variation at neutral loci is unlikely to reflect adaptive genetic diversity (Bonin, Nicole, Pompanon, Miaud, & Taberlet, 2007; Holderegger, Kamm, & Gugerli, 2006; McKay & Latta, 2002; Teixeira & Huber, 2021). *Ex situ* population sampling may need to evaluate the impact different evolutionary processes have had on population genetic structure to optimize neutral and adaptive variation collected. Thus, an understanding of population connectivity and the impact of selection across populations can inform *ex situ* collection design. If empirical or simulated data suggest populations are genetically connected and genetic differentiation is low, then most neutral genetic variation may be captured within one or a few populations. However, if selection overcomes the homogenizing effects of gene flow, ensuring adaptive genetic differences are preserved for all populations will require the inclusion of diverse population origins, separation of such populations *ex situ*, and consideration of population origin in breeding programs.

With the advent of next-generation sequencing (NGS) and the ever-decreasing costs associated with these technologies, genome-wide estimates of genetic diversity can be readily assessed and used to guide conservation management strategies. Combined with statistical and simulation-based tools, these data provide a powerful and timely means to evaluate aspects of population genetic variation and spatial genetic structure critical to informing genetic rescue and *ex situ* conservation plans, including both populations’ demographic and adaptive history (Abebe, Naz, & Léon, 2015; Liu, Zhang, Wang, & Ma, 2020; Wang, Bernhardsson, & Ingvarsson, 2020; Xia et al., 2018). However, in conifers, despite extensive use in characterizing genomes as well as neutral and adaptive variation (Eckert et al., 2010; Namroud, Beaulieu, Juge, Laroche, & Bousquet, 2008; Nystedt et al., 2013; Stevens et al., 2016; Tyrmi et al., 2020; Wang et al., 2020), these data have only rarely been used to inform conservation management decisions.

Torrey pine (*Pinus torreyana* Parry) is a critically endangered pine (IUCN, 2021), endemic to California. One of the rarest pine species in the world (Critchfield & Little, 1966; Dusek, 1985), Torrey pine’s distribution spans one island population (*Pinus torreyana* subsp. *insularis*) of approximately 3,000 reproductive individuals (Santa Rosa Island, CA), and one mainland population (*Pinus torreyana* subsp. *torreyana*) of approximately 4,000 reproductive individuals (Torrey Pine State Reserve in La Jolla, CA) (J. Franklin & Santos, 2011; Hall & Brinkman, 2015). In addition to low population size, and despite current *in situ* and *ex situ* conservation efforts, Torrey pine suffers from exceedingly low genetic variation and faces both anthropogenic and environmental disturbances (J. Franklin & Santos, 2011; Hamilton, Royauté, Wright, Hodgskiss, & Ledig, 2017; Ledig & Conkle, 1983; Waters & Schaal, 1991; Whittall et al., 2010). For these reasons, the species may be at imminent risk for population-scale extirpation events and thus a potential candidate for genetic rescue. Inter-population admixture may increase population genetic diversity, alleviating potential fitness consequences associated with Torrey pine’s low genetic diversity (Hamilton et al., 2017), and increase evolutionary potential necessary to respond to current and future ecological challenges (Carlson, Cunningham, & Westley, 2014). Previous research observed heterosis following one generation of admixture between island and mainland individuals, suggesting that genetic rescue may alleviate fitness consequences associated with reduce genetic variation (Hamilton et al., 2017). However, if adaptive genetic differences have evolved between island and mainland populations, fitness consequences following the disruption of co-adapted gene complexes may not be observed in the first generation cross. Thus, although the combination of exceedingly low genetic diversity and conservation status suggest Torrey pine may be a candidate for genetic rescue, evaluation of the species’ demographic and adaptive evolutionary history will be necessary prior to inform conservation management decisions.

With this study, we use genomic data to quantify and model aspects of populations’ demographic and evolutionary history necessary to preserve rare species’ evolutionary potential. Specifically, we delineate the contribution of stochastic and deterministic processes to genomic differentiation in *Pinus torreyana*, asking three questions: (i) what are current effective population sizes and how have they changed over time, (ii) have populations remained genetically connected following isolation, and (iii) is there evidence of adaptive divergence between populations that may indicate distinct evolutionary trajectories? We identify the most probable demographic scenario for Torrey pine using Approximate Bayesian Computing (ABC) and coalescent simulations. To evaluate the role of selection, we identify loci that may be important to adaptation using various F_ST_ outlier methods and assess the functional significance of these loci by annotating candidates with the gene ontology (GO) resource. Using this knowledge, we discuss conservation strategies for Torrey pine, including *ex situ* sampling to preserve neutral and nonneutral processes within species collections, and possible risks associated with genetic rescue. This study demonstrates the benefits and necessity of understanding the demographic and adaptive history of rare species to guide conservation.

## Material and Methods

### Population sampling and DNA extraction

Between June and July 2017, needle tissue was collected from individuals spanning the entire natural distribution of *Pinus torreyana* (Torrey pine; Fig. 1). A total of 286 individuals were sampled, including 146 individuals from the mainland population at the Torrey Pine State Reserve (TPSR) near La Jolla, CA and 140 individuals from the island population on Santa Rosa Island, CA (SRI), one of the Channel Islands. Needles were dried in silica gel following which genomic DNA was extracted using between 25-30 mg of dry needle tissue and a modified CTAB protocol (Doyle & Doyle, 1987). To reduce DNA shearing, slow manual shaking of tubes was used. Following extraction, the concentration and purity of DNA extracted was quantified for each sample using a NanoDrop 1000 Spectrophotometer (Thermo scientific) to ensure all samples had a concentration of approximately 85 ng/µl and purity ratios of 1.4 and above (average across samples; 260/280 = 1.85, 260/230 = 1.96).

**Figure 1.**
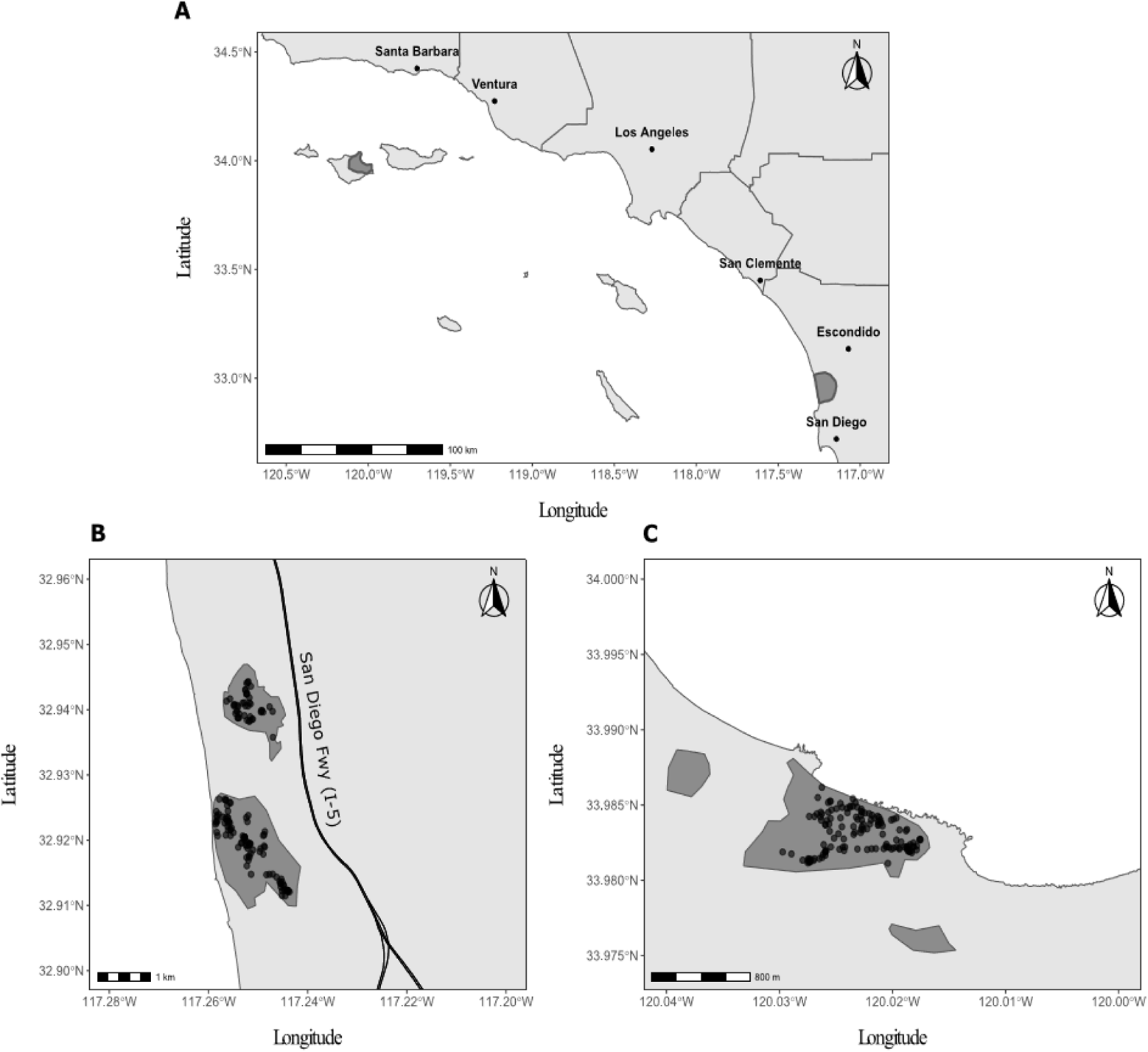
(A) Distribution of the last two remanent populations of Torrey pine (dark grey shades). Top left: Santa Rosa Island, Channel Islands, CA. Bottom right: Torrey Pine State Reserve, La Jolla, CA. (B-C) Population-specific distribution of Torrey pine (dark grey shades) and trees sampled for needle tissue (circles) at the Torrey Pine State Reserve (TPSR, B) and on Santa Rosa Island (SRI, C).

### Genomic library preparation and ddRAD sequencing

Genomic libraries were prepared for all 286 individuals following the protocol of Parchman & colleagues (2012). Briefly, 510 ng (6 µl at 85 ng/µl) of DNA was digested using endonucleases *Eco*RI and *Mse*I (New England BioLabs, Inc.) after which barcoded (*Eco*RI cut site) and non-barcoded (*Mse*I cut site) adapters compatible with Illumina sequencing were ligated to each end of DNA fragments using T4 ligase (New England BioLabs, Inc.). A different barcode sequence was used for each of the 286 samples. Due to the large, highly repetitive nature of pines’ genome (Stevens et al., 2016), we used the methylation-sensitive enzyme *Eco*RI, as it effectively reduces the presence of repetitive and non-coding DNA sequences in genomic libraries (Parchman et al., 2012). Restriction-ligation products were amplified using two successive PCR reactions to produce genomic libraries with concentrations necessary for sequencing (> 2nM). All PCR-amplified genomic libraries were subsequently pooled and sent to the Genomic Sequencing and Analysis Facility (GSAF; Austin, TX) for size selection of fragments within the range of 450-500 bp and sequenced on 5 lanes of an Illumina HiSeq 2500 using the 100 bp single-end sequencing protocol (1 × 100 bp).

### De novo assembly and SNP calling

Demultiplexing of sequence files was performed using *ipyrad* v0.9.12 (Eaton & Overcast, 2020) allowing one mismatch in the barcode sequence. Reads were filtered, assembled *de novo*, and used to call SNPs within the *dDocent* v2.7.8 pipeline (Puritz, Hollenbeck, & Gold, 2014; Puritz, Matz, et al., 2014). Reads were filtered by removing low-quality bases at the beginning and end of reads (PHRED score < 20), Illumina adapters, and trimmed when the average PHRED score fell below 10 within a 5 bp window using the program TRIMMOMATIC (Bolger, Lohse, & Usadel, 2014). As a contiguous genome assembly for *P. torreyana* is not available, we first generated a reference of genomic regions sampled with our sequencing design using a *de novo* approach. Reads were clustered based on sequence similarity and assembled into a reference assembly using the program CD-HIT (Fu, Niu, Zhu, Wu, & Li, 2012; W. Li & Godzik, 2006). To be included in *de novo* assembly, reads had to have a minimum of 3x within-individual coverage and be present in at least 5 individuals. To form a cluster (locus), reads had to have a minimum of 86% sequence similarity, a cutoff previously used in pines (Menon et al., 2018). This threshold was chosen as a tradeoff to avoid the clustering of paralogous loci while still accounting for the presence of missing bases, errors, or polymorphisms between true homologous sequences. Finally, reads were mapped onto *de novo* assembled loci using BWA MEM (H. Li, 2013) and SNPs were called using the software FREEBAYES (Garrison & Marth, 2012). Read mapping was performed using BWA default parameters, including a match value of 1, a mismatch penalty of 4 and a gap penalty of 6. This yielded a set of 652,492 SNPs that were subjected to downstream filtering. Variants with genotype quality (GQ) < 20 and genotype depth < 3 were first marked as missing. Then, variants with PHRED scores (QUAL) ≤ 30, minor allele counts < 3, minor allele frequencies < 0.01, call rate across all individuals < 0.95, mean depth across samples > 57 (based on the equation: *d* + 4√*d*, where *d* is the average read depth across variants, H. Li, (2014), F_IS_ estimates < −0.5 or > 0.5, and linkage score (r^2^) > 0.5 within a 95 bp window were removed from the raw SNP dataset. Following filtering, a total of 93,085 biallelic SNPs were kept and used for analysis (hereafter referred to as the full dataset). Note that 16 individuals with > 40% missing data were also discarded, leaving 270 genotyped individuals (SRI: 130 individuals, TPSR: 140 individuals) for inclusion in analyses.

### Population structure and genetic diversity analyses

To describe and quantify contemporary genetic differences between Torrey pine populations, we first assessed genetic structure of populations using principal component analysis (PCA) implemented in the R package ADEGENET (Jombart, 2008; Jombart & Ahmed, 2011). Unless otherwise stated, analyses were performed in R version 3.6.2 and 4.0.2 (R Core Team, 2019, 2020). To quantify genetic differences between island and mainland populations, we calculated Nei’s F_ST_ statistic (Nei, 1987) for each SNP using the HIERFSTAT R package (Goudet & Jombart, 2020) and averaged estimates across loci. A 95% confidence interval around the mean was constructed by bootstrapping the empirical F_ST_ distribution 10,000 times using R packages BOOT (Canty & Ripley, 2021; Davison & Hinkley, 1997) and SIMPLEBOOT (Peng, 2019).

To determine the extent of contemporary within-population genetic structure, the most likely number of genetic clusters was independently evaluated for SRI and TPSR using the function *find.clusters()* implemented in the R package ADEGENET. This function transforms genomic data using principal component analysis and performs successive K-means clustering with an increasing number of clusters (*k*). For each successive value of *k*, the Bayesian Information Criterion (BIC) is computed and was used to assess the optimal number of clusters (*k*). For each population, we assessed between 1 to 10 clusters while retaining principal components necessary to explain 90% of the variation after ordination (SRI: 114, TPSR: 122). For TPSR, two individuals (TPSR5107, TPSR3189) clustered distantly from the population, which may mask subtle within-population genetic structure. Consequently, we re-ran the analysis excluding the two individuals while maintaining the same range for *k* (1 to 10) and retaining 121 principal components (90.38% of variation explained after ordination).

To evaluate Torrey pine evolutionary potential, contemporary standing genetic variation within the species was estimated by calculating expected heterozygosity (H_E_), inbreeding coefficients (F_IS_), and coancestry coefficients (***θ***) for both island and mainland populations independently. Values of H_E_ and F_IS_ were calculated for each SNP separately using the R package ADEGENET and averaged across loci to provide population-level estimates. To evaluate ***θ***, we used the R package RELATED (Pew, Wang, Muir, & Frasier, 2015) that estimates genetic relatedness between all possible pairs of individuals within a population. Specifically, we used the triadic likelihood (TrioML) estimate of relatedness assuming no inbreeding within populations. Averaging pairwise ***θ*** values across all individuals within TPSR and SRI provided population estimates of genetic relatedness. To build 95% confidence intervals around H_E_, F_IS_ and ***θ*** averages, the empirical distribution for each parameter within each population was bootstrapped 10,000 times in R using the BOOT and SIMPLEBOOT packages.

### ABC demographic modeling

***Demographic models –*** To evaluate the impact genetic drift and gene flow have had to patterns of neutral genetic variation in Torrey pine, we quantified changes in effective population size over time, interpopulation migration rate, and time since population divergence. To do so, we tested six distinct demographic models that were classified into two broad categories: (1) isolation with/without migration (Fig. 2A), and (2) two-population demic expansion (Fig. 2B, C).

**Figure 2.**
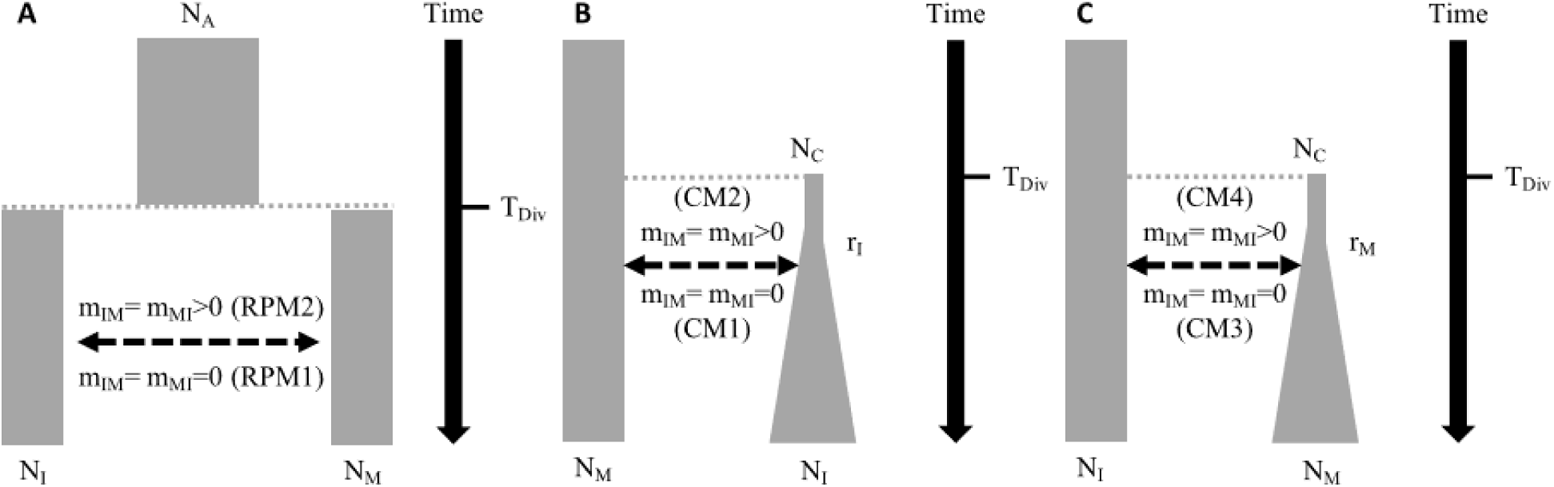
Schematics of demographic scenarios simulated. Rectangles represent current or ancestral populations, dashed arrows represent migration between population, and solid arrow represent time. (A) Scenarios of isolation without (RPM1) or with (RPM2) migration between populations. (B) Island colonization scenarios without (CM1) or with (CM2) subsequent migration between populations. (C) Mainland colonization scenarios without (CM3) or with (CM4) subsequent migration between populations. N_A_, ancestral effective population size; N_I_, island effective population size; N_M_, mainland effective population size; N_C_ initial effective population size following migration (number of migrants); m_IM_, migration probability from island to mainland; m_MI_, migration probability from mainland to island; T_Div_, time of population divergence; r_I_, island (exponential) growth rate; r_M_, mainland (exponential) growth rate

Models of isolation with or without migration (RPM1, RPM2) were developed to test a hypothesis formulated by Ledig & Conkle, (1983). This hypothesis predicts that there was once a single ancestral population of Torrey pine that diverged to form one island and one mainland population following tectonic movement (Fig. 2A). These models assume that an ancestral population with an effective size N_A_ diverged T_Div_ generations before present to form two populations with current effective sizes N_I_ (island, SRI) and N_M_ (mainland, TPSR). Following divergence, to assess whether gene flow has occurred between populations, bidirectional migration was either prevented (RPM1, m_IM_=m_MI_=0) or permitted (RPM2, m_IM_=m_MI_>0). Both models assume constant island and mainland effective population size following divergence.

The remaining four models (CM1, CM2, CM3, CM4) tested two different hypotheses of land colonization where one population was founded by the other (Fig. 2B, C). Models CM1 and CM2 specifically test the hypothesis that Santa Rosa Island was colonized by a subset of mainland individuals (Ledig & Conkle, 1983). In this scenario, SRI was founded by N_C_ effective migrants from TPSR T_Div_ generations ago and grew exponentially at a rate 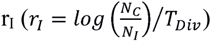 to form a population with an effective size N_I_. TPSR effective population size (N_M_) was assumed constant. As above, bidirectional migration between populations was either prevented (CM1, m_IM_=m_MI_=0) or permitted (CM2, m_IM_=m_MI_>0) to evaluate whether gene flow has occurred between populations following colonization. Models CM3 and CM4 test the hypothesis that the mainland population was founded by a subset of island individuals (Haller, 1986). This scenario assumes TPSR was founded by N_C_ effective migrants from SRI T_Div_ generations before present. SRI effective population size (N_I_) was assumed constant while TPSR effective population size (N_M_) grew exponentially at 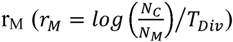. Once again, to test whether gene flow has occurred between populations since colonization, exchange of migrants between population was either prevented (CM3, m_IM_=m_MI_=0) or permitted (CM4, m_IM_=m_MI_>0).

For all six models, uniform priors were used except for T_Div_, N_C_, and N_A_ for which we used log-uniform priors. Priors on a logarithmic scale increases the weight given to small values and is recommended when parameters’ ranges span several orders of magnitude (Wegmann, Leuenberger, & Excoffier, 2009). Details on demographic parameters and their prior distributions are provided in Table 1.

**Table 1.**
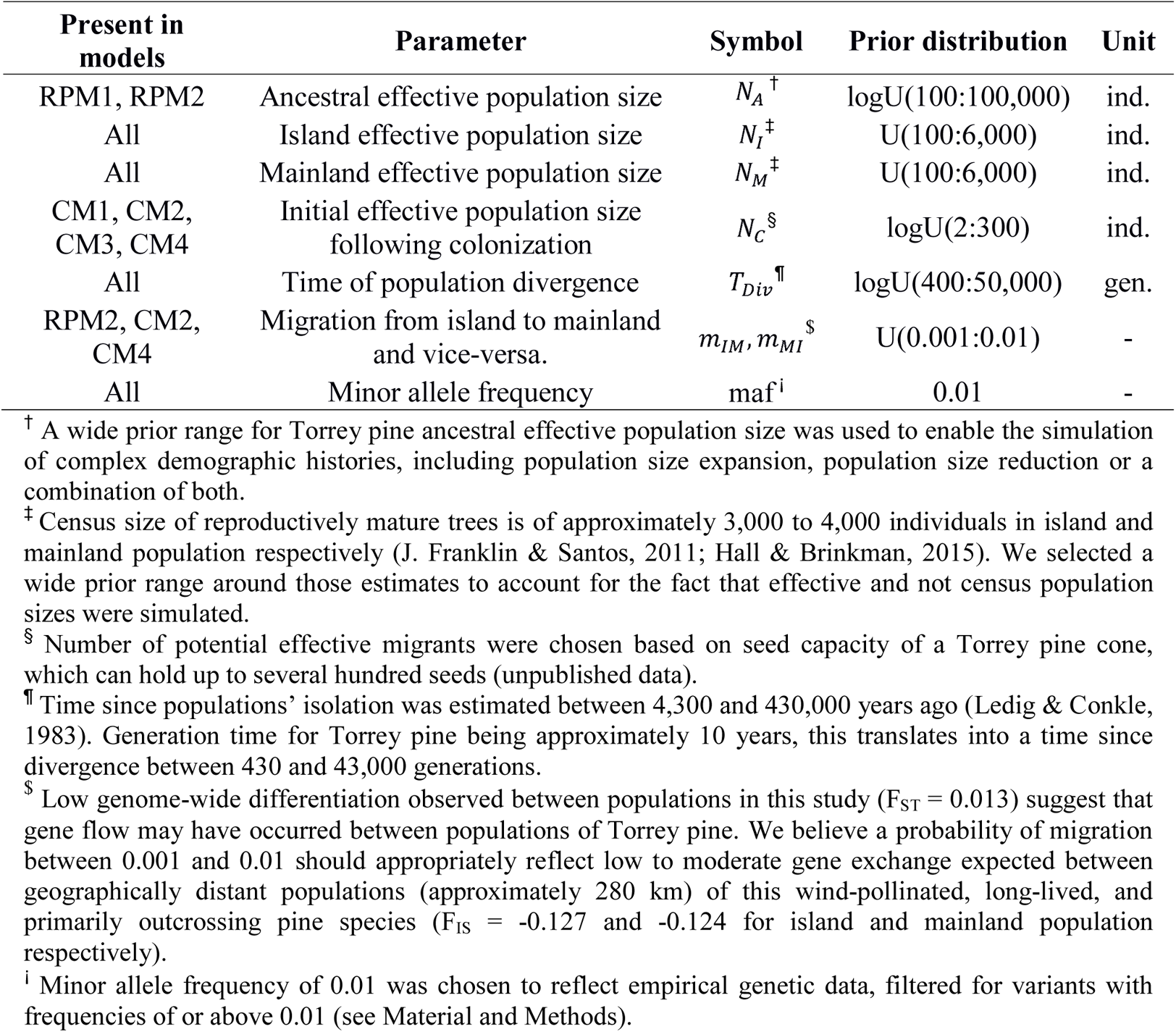
Demographic parameters with their prior distributions and occurrence in each of the six models simulated.

| Present in models | Parameter | Symbol | Prior distribution | Unit |
| --- | --- | --- | --- | --- |
| RPM1, RPM2 | Ancestral effective population size | $N_A^{\dagger}$ | logU(100:100,000) | ind. |
| All | Island effective population size | $N_I^{\ddagger}$ | U(100:6,000) | ind. |
| All | Mainland effective population size | $N_M^{\ddagger}$ | U(100:6,000) | ind. |
| CM1, CM2, CM3, CM4 | Initial effective population size following colonization | $N_C^{\S}$ | logU(2:300) | ind. |
| All | Time of population divergence | $T_{Div}^{\P}$ | logU(400:50,000) | gen. |
| RPM2, CM2, CM4 | Migration from island to mainland and vice-versa. | $m_{IM}, m_{MI}^{\$}$ | U(0.001:0.01) | - |
| All | Minor allele frequency | maf<sup>i</sup> | 0.01 | - |

<sup>†</sup> A wide prior range for Torrey pine ancestral effective population size was used to enable the simulation of complex demographic histories, including population size expansion, population size reduction or a combination of both.
<sup>‡</sup> Census size of reproductively mature trees is of approximately 3,000 to 4,000 individuals in island and mainland population respectively (J. Franklin & Santos, 2011; Hall & Brinkman, 2015). We selected a wide prior range around those estimates to account for the fact that effective and not census population sizes were simulated.
<sup>§</sup> Number of potential effective migrants were chosen based on seed capacity of a Torrey pine cone, which can hold up to several hundred seeds (unpublished data).
<sup>¶</sup> Time since populations' isolation was estimated between 4,300 and 430,000 years ago (Ledig & Conkle, 1983). Generation time for Torrey pine being approximately 10 years, this translates into a time since divergence between 430 and 43,000 generations.
<sup>\$</sup> Low genome-wide differentiation observed between populations in this study ( $F_{ST} = 0.013$ ) suggest that gene flow may have occurred between populations of Torrey pine. We believe a probability of migration between 0.001 and 0.01 should appropriately reflect low to moderate gene exchange expected between geographically distant populations (approximately 280 km) of this wind-pollinated, long-lived, and primarily outcrossing pine species ( $F_{IS} = -0.127$ and $-0.124$ for island and mainland population respectively).
<sup>i</sup> Minor allele frequency of 0.01 was chosen to reflect empirical genetic data, filtered for variants with frequencies of or above 0.01 (see Material and Methods).

***Additional filtering and down-sampling of genetic variants –*** Accurate estimation of demographic parameters using coalescence simulations requires the use of neutrally evolving genetic markers. To ensure neutrality of SNPs, we filtered the full dataset for SNPs that did not deviate significantly from Hardy-Weinberg equilibrium (HWE). Since population structure may create departures from HWE, we applied this filter to island and mainland populations independently and removed SNPs that deviated significantly from HWE (P < 0.05) in both populations. This was performed using a customized R function relying on R packages ADEGENET and PEGAS (Paradis, 2010). In total, 73,928 SNPs were retained following filtering for use in demographic modelling (hereafter referred to as the HWE-filtered dataset).

For computational efficiency, we down sampled the HWE-filtered dataset from 73,928 to 9,795 variants first by generating bivariate bins based on observed heterozygosity and Nei’s F_ST_ (0.05-interval bins), and then by subsampling each bin proportionally to the number of SNPs they contained. In this way, each bin is subsampled to reflect its contribution to the HWE-filtered dataset (Appendix S1). Following subsampling, we conducted a principal component analysis using the down-sampled dataset to ensure patterns of genetic diversity and population structure were maintained between datasets (Appendix S2).

***Generating coalescent simulations and estimating summary statistics –*** To evaluate and compare demographic scenarios, a set of 200,000 simulations was generated using ABCSAMPLER for each model, a wrapper program included in the package ABCTOOLBOX (Wegmann, Leuenberger, Neuenschwander, & Excoffier, 2010). For each simulation, ABCSAMPLER samples prior ranges of demographic parameters and uses these values as inputs for coalescence simulations within a user-defined simulation program. We used FASTSIMCOAL version 2.6.0.3 (Excoffier, Dupanloup, Huerta-Sánchez, Sousa, & Foll, 2013; Excoffier & Foll, 2011) to simulate 9,795 unlinked SNPs with a minor allele frequency of 0.01, reflecting the composition of the down-sampled genetic dataset. For each model, simulated data were output as genotypes and fed to a user-defined program by ABCSAMPLER to estimate population genetic summary statistics. ARLEQUIN version 3.5.2.2 (Excoffier & Lischer, 2010) was used to calculate ten distinct population genetic summary statistics, specifically aiming at quantifying genetic variation and divergence within and between Torrey pine populations. These included genetic diversity (i.e., population-specific heterozygosity and number of alleles, average heterozygosity and number of alleles over loci and populations, and mean total heterozygosity), genetic differentiation (i.e., pairwise F_ST_), and variance (i.e., standard deviation over populations of the average heterozygosity and number of alleles) statistics. Finally, to obtain observed population genetic summary statistics, ARLEQUIN version 3.5.2.2 was rerun using the down-sampled dataset. Characterizing and summarizing the amount and distribution of genetic variation within each dataset, these statistics were used to calculate posterior probabilities and identify the demographic model with greatest support, as well as estimate demographic parameters associated with that model (see below).

***ABC parameter estimation –*** Demographic parameters were estimated in R using the ABC package (Csilléry, François, & Blum, 2012). Cross-validation simulations were conducted first to evaluate the ability of summary statistics to distinguish between demographic models (Appendix S3). We performed leave-one-out cross-validations, consisting of selecting one simulation of a demographic scenario and then assigning it to one of the six models using posterior probabilities estimated from all remaining simulations. This was repeated one hundred times for each demographic model, generating a confusion matrix. If misclassification rates (proportions of simulations incorrectly assigned to a model) are low, then computed summary statistics can distinguish between our different demographic scenarios. The posterior probability of each demographic model was approximated as the proportion of accepted simulations and used to select the best model following cross-validation. To ensure the best model provided a good fit to the data, we performed a goodness-of-fit test as implemented in the function *gfit()*. Finally, demographic parameters associated with the best model were estimated as the weighted medians of posterior distributions using the *weighted.median()* function implemented in the R package SPATSTAT (Baddeley, Rubak, & Turner, 2015). Posterior distributions were created from the set of accepted simulations using a non-linear postsampling regression adjustment conducted on log-transformed data (Blum & François, 2010). Ninety-five percent confidence intervals around weighted medians were estimated using 10,000 bootstrap replicates of posterior distributions in R, using BOOT, SIMPLEBOOT and SPATSTAT packages. The validity and accuracy of each estimated parameter were tested using additional, yet distinct, 100-fold leave-one-out cross-validation simulations (Appendix S4). The cross-validation process begins with the random selection of one simulation generated by the best demographic model. Summary statistics associated with that simulation are considered as pseudo-observed data and its demographic parameters are estimated using remaining simulations for the model. If pseudo-observed parameters can accurately be predicted, then inferred demographic parameters from true observed data can be considered valid and accurate. Cross-validation simulations, model selection, model validation, and parameters estimation were conducted using a tolerance threshold of 0.01, a tradeoff between retaining a reasonable number of simulations to estimate posterior distributions and keeping the tolerance value as low as possible (S. Li & Jakobsson, 2012). Finally, as census sizes for Torrey pine populations are available (J. Franklin & Santos, 2011; Hall & Brinkman, 2015), we calculated the proportion of the census size (N) to effective population size (Ne) for each population separately. Of all reproductive trees present within a population (census size), this ratio estimated the proportion contributing to the next generation (effective size).

### Simulating the null F_ST_ distribution

To evaluate the influence natural selection may have had on the genomic structure of Torrey pine populations, we compared the distribution of Nei’s F_ST_ estimated from the full SNP dataset with a simulated distribution based on 93,085 independent SNPs from 270 individuals generated using SIMCOAL2 version 12.09.07 (Laval & Excoffier, 2004). We used weighted medians estimated from posterior distributions borrowed from the best demographic model (see Results) as input parameters for neutral simulations. For each simulated SNP, the minimum allele frequency was set to 0.01 to reflect filters applied to the full dataset. Locus-specific Nei’s F_ST_ values were estimated for both full and simulated datasets in R using the HIERFSTAT package (Appendix S5).

### Outlier detection analyses

To estimate the potential contribution of local adaptation to genomic differentiation between Torrey pine populations, we used the full dataset of 93,085 SNPs to identify loci putatively under selection using three distinct methods: BAYESCAN version 2.1 (Foll & Gaggiotti, 2008), OUTFLANK version 0.2 (Whitlock & Lotterhos, 2014) and PCADAPT version 4.3.3 (Privé, Luu, Vilhjálmsson, & Blum, 2020). These three approaches were selected for their ability to account for neutral population structure (BAYESCAN, OUTFLANK) and to handle genetic admixture between individuals (PCADAPT). For both BAYESCAN and OUTFLANK, we grouped all 270 individuals by populations (mainland: 140 individuals, island: 130 individuals). Below is a brief description of all three methods used to identify candidate SNPs.

The first approach we used was BAYESCAN. For each locus, BAYESCAN uses a Bayesian approach to decompose F_ST_ coefficients into population- and locus-specific components using a logistic regression. Loci are identified as putatively under selection if the locus-specific component is needed to explain the observed distribution of genetic diversity. Our analysis was conducted using BAYESCAN default parameters. Results were visualized and analyzed in R. Only SNPs with a false discovery rate of 1% or below were retained and considered as candidate loci. The second approach we used was OUTFLANK, an R package which identifies outliers by inferring a null F_ST_ distribution approximated from empirical F_ST_ values. This distribution was produced by removing loci with an expected heterozygosity below 0.1 (Hmin = 0.1), trimming 5% of lowest and highest empirical F_ST_ values (RightTrimFraction = LeftTrimFraction = 0.05), and retaining only loci passing a built-in neutrality test with a false discovery rate of 0.1% (qthreshold = 0.001). Using the latter threshold provided a conservative estimate for neutral F_ST_. Outlier loci were identified by comparing the empirical F_ST_ distribution to the inferred null F_ST_ distribution using a built-in chi-squared test with a false discovery rate of 1% (qthreshold = 0.01). Note that loci with H_E_ < 0.1 were excluded while conducting the chi-squared test and therefore could not be identified as potential outliers (Hmin = 0.1). The third and last approach we used was PCADAPT. Also implemented in R, PCADAPT is a package that assesses population structure using PCA. Consequently, this approach does not require individuals to be grouped into populations. Following PCA, candidate loci are identified as those substantially correlating with population structure. We ran PCADAPT retaining the first axis of differentiation (K = 1) and SNPs with a minor allele frequency of 0.01 (min.maf = 0.01). We only considered the first principal component to calculate the test statistic, as additional axes did not ascertain population structure (Appendix S6). Candidate loci were identified as the set of SNPs with a false discovery rate of 1% or below. To minimize the presence of false positives within our dataset, only outlier SNPs common to all three approaches were considered as candidate loci and included in subsequent analyses. Finally, to both visualize and quantify genetic structure at putatively adaptive loci, we conducted a principal component analysis based only on candidate SNPs shared by all three methods using the R package ADEGENET.

### Functional categorization of candidate loci

To identify biological processes or molecular functions that may play a role in adaptation to mainland or island environments, *de novo* assembled sequences containing candidate SNPs common to all genome scans were first extracted and then annotated using BLASTx version 2.6.0+. Sequences were blasted against the UniProt protein database filtered for sequences from species within the *Pinaceae* family (Taxon identifier 3318). Sequence similarity was assessed using BLASTx default parameters, an e-value hit filter of 10^−3^ (−evalue 0.001), and a number of database hits to retain of 1 (-max_target_seqs 1). Gene ontology terms were mapped onto annotated sequences in R using the UNIPROTR package (Soudy et al., 2020).

## Results

### Population genetic structure and variation

Using all 93,085 SNP markers, the principal component analysis revealed little genome-wide differentiation in Torrey pine with the first two principal components (PC1, PC2) explaining only 1.9 and 0.6% of observed genetic differences, respectively (Fig. 3). Nonetheless, PC1 unambiguously separated island from mainland individuals, suggesting some level of genetic differentiation exists between populations. These results were further supported by the low average coefficient of genetic differentiation found across loci (Nei’s F_ST_), estimated at 0.013 (95% CI: 0.012 - 0.013). In addition to population-scale genetic differentiation, within-population genetic differentiation was estimated to evaluate whether local inbreeding, possibly increased in small populations, could have contributed to fine-scale genetic structure in Torrey pine. For both island and mainland populations, we found no evidence of within-population genetic structure. Population-specific principal component analyses identified no clear genetic clusters and revealed that, combined, the first two axes of differentiation (PC1, PC2) only explained approximately 1.9% and 2.2% of mainland and island within-population genetic differences, respectively (Fig. 4). In addition, evaluating the likelihood between 1 to 10 genetic clusters (*k*) within each population using BIC indicated that populations appear largely homogeneous (most likely *k* = 1) (Appendix S7). Interestingly, average estimates of inbreeding and coancestry coefficients across loci were low for both the island and the mainland population (Table 2). While low inbreeding coefficients support the lack of observable within-population genetic structure, this also indicates that reproduction among relatives or unequal reproductive success among individuals is unlikely to have contributed to low expected heterozygosity observed within populations (Table 2). Combined, our results indicate that Torrey pine exhibits exceedingly low genetic diversity, with the majority of variation distributed within genetically unstructured populations.

**Figure 3.**
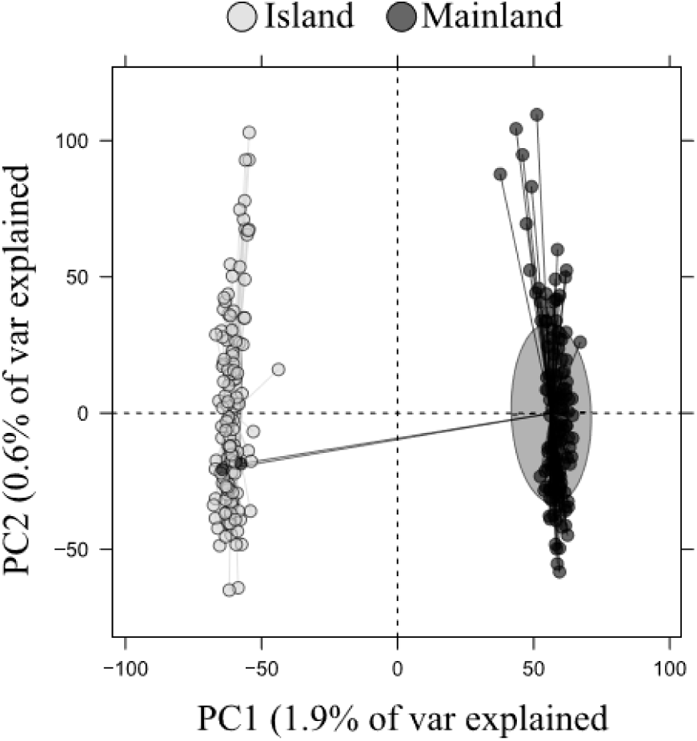
Principal component analysis using 93,085 SNPs for 270 Torrey pine individuals, including individuals from both mainland (black) and island (grey) populations. Variation explained by the first two principal components is provided in parentheses.

**Figure 4.**
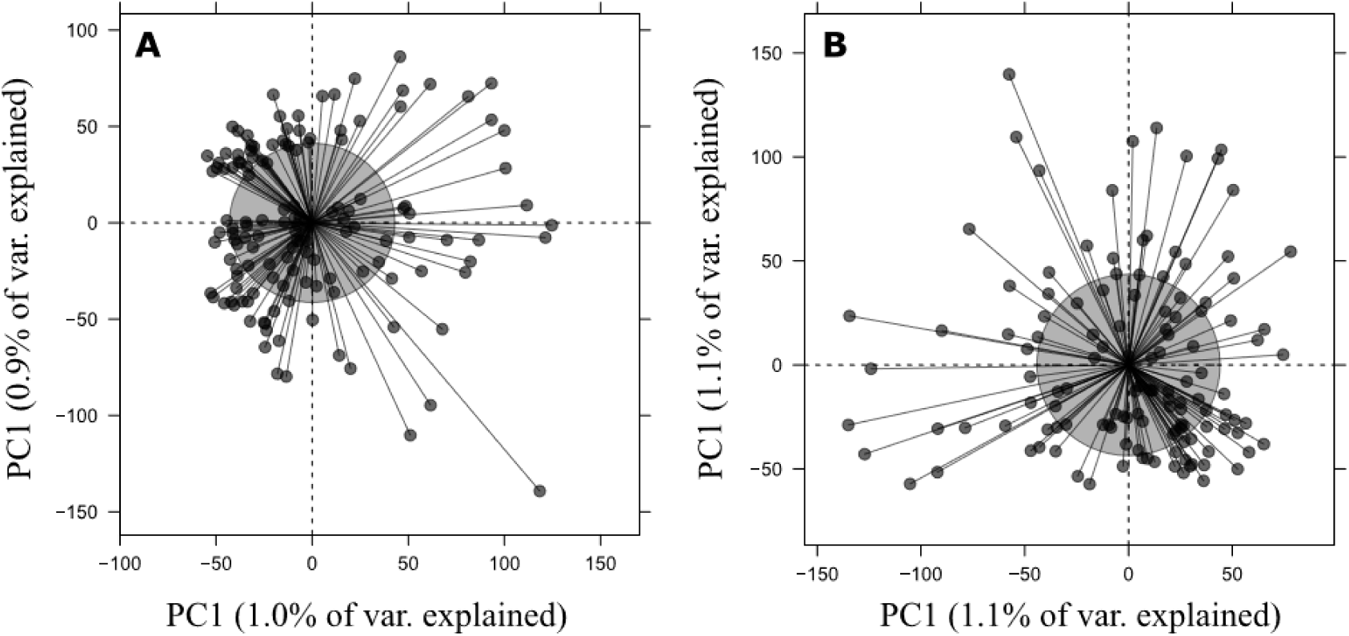
Population-specific principal component analysis based on 93,085 SNP variants. (A) Mainland population (TPSR, n = 138). Note that two individuals were removed from analysis to visualize within-population structure on a finer scale (see Material and Methods). (B) Island population (SRI, n = 130).

**Table 2.** Average genetic summary statistics across loci, including expected heterozygosity (H_E_), inbreeding coefficient (F_IS_), and coancestry (relatedness) coefficient (*θ*) for island (SRI) and mainland (TPSR) Torrey pine populations. Ninety-five percent confidence interval (95% CI) around mean estimates were obtained by bootstrapping.

| Population | $H_E$<br>(95% CI) | $F_{IS}$<br>(95% CI) | $\theta$<br>(95% CI) |
| --- | --- | --- | --- |
| Island | 0.185<br>(0.184, 0.186) | -0.127<br>(-0.128, -0.126) | 0.023<br>(0.023, 0.024) |
| Mainland | 0.184<br>(0.183, 0.184) | -0.124<br>(-0.125, -0.124) | 0.022<br>(0.022, 0.023) |

### Demographic history of Torrey pine

Of the six demographic models evaluated (Fig. 2), the isolation with migration model (RPM2) received the most support with a posterior probability of 92.18%. The remaining five models exhibited lower posterior probabilities ranging from 0% to 3.98%. With low misclassification rates, cross-validations indicated that simulated summary statistics were able to confidently distinguish between different demographic scenarios (Appendix S3). The goodness-of-fit test revealed that simulated summary statistics did not significantly differ from observed ones (P = 0.76), providing a good fit to the data.

Based on RPM2, an ancestral Torrey pine population with an effective size of approximately 1,124 individuals (95% CI: 939 - 1,213) diverged during the early Pleistocene approximately 1.2 million YBP (95% CI: 1,195,367 - 1,296,186, assuming a generation time of ten years) to form one island and one mainland population with effective sizes of approximately 2,305 (95% CI: 2,166 - 2,338) and 1,715 (95% CI: 1,616 - 1,759) individuals, respectively (Fig. 5). This resulted in a 0.75 (N_I_/N = 2,305/3,063) proportion of the census to effective population size on the island, and a 0.45 (N_M_/N = 1,715/3,806) proportion of the census to effective population size on the mainland. Following divergence, some gene flow was maintained between populations with an estimated migration rate of 8.34 × 10^−3^ (95% CI: 8.15 × 10^−3^ - 8.76 × 10^−3^) per generation. In general, cross-validation simulations indicated low prediction errors (Appendix S4), suggesting high accuracy of inferred parameters. Nonetheless, note that for T_Div_, the associated prediction error was higher than for other parameters.

**Figure 5.**
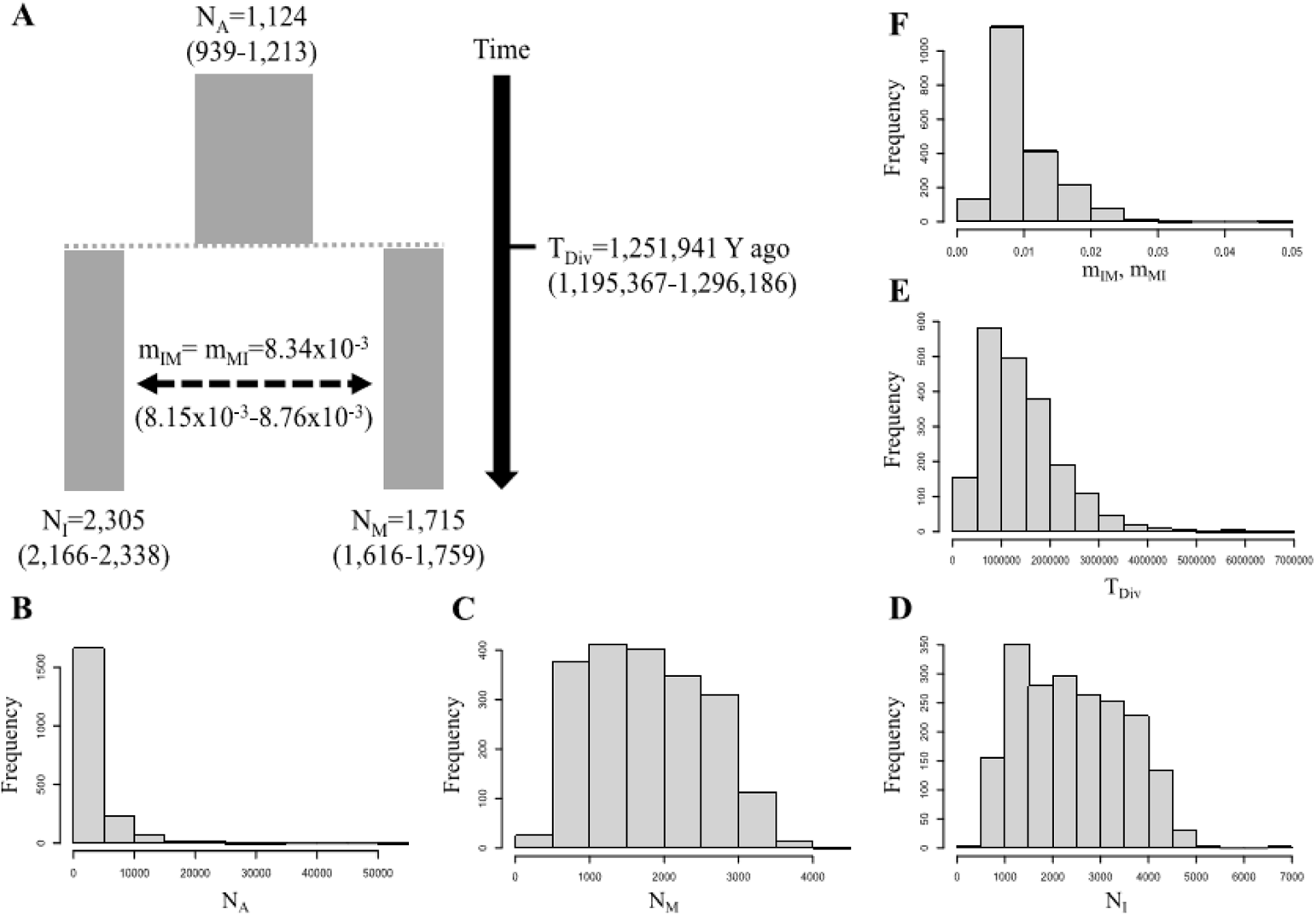
(A) Demographic parameters (weighted medians) and 95% confidence intervals (in parentheses) estimated using RPM2. Each rectangle represents a population, either contemporary or ancestral. The solid arrow represents time, while the dashed arrow indicates gene flow between populations. (B-F) Posterior distribution of each demographic parameter inferred using a tolerance rate of 0.01. N_A_, ancestral effective population size; N_I_, island effective population size; N_M_, mainland effective population size; m_IM_, migration probability from island to mainland; m_MI_, migration probability from mainland to island; T_Div_, time of population divergence.

### Divergent selection between island and mainland populations

Despite limited genetic variation within populations, we found some evidence for the evolution of genetic differences among populations at a subset of loci. We compared the neutral F_ST_ distribution generated from the simulated RPM2 demographic model with the empirical distribution based on our full dataset of SNP variants (Appendix S5). For select SNPs, moderate to high empirical F_ST_ values (from approximately 0.2 to 0.65) could not be generated through neutral simulations, suggesting they may be candidate for selection. PCADAPT, BAYESCAN, and OUTFLANK identified 3,138 (3.37%), 2,163 (2.32%), and 3,942 (4.23%) outlier SNPs, respectively (Fig. 6A). Of these outlier loci, 2,053 (2.21%) were common to all three methods and contribute to genomic structure between Torrey pine populations (Fig. 6B). Indeed, the principal component analysis revealed that the first axis of differentiation (PC1) unambiguously differentiated island from mainland populations based on common outlier SNPs, explaining over 20% of observed genetic differences. Consistent with these results, F_ST_ values for putatively adaptive loci ranged between 0.1 and 0.63 and either could not be generated or could only be generated at low frequency through neutral simulations, representing only 0.078% of all simulated F_ST_ values.

**Figure 6.**
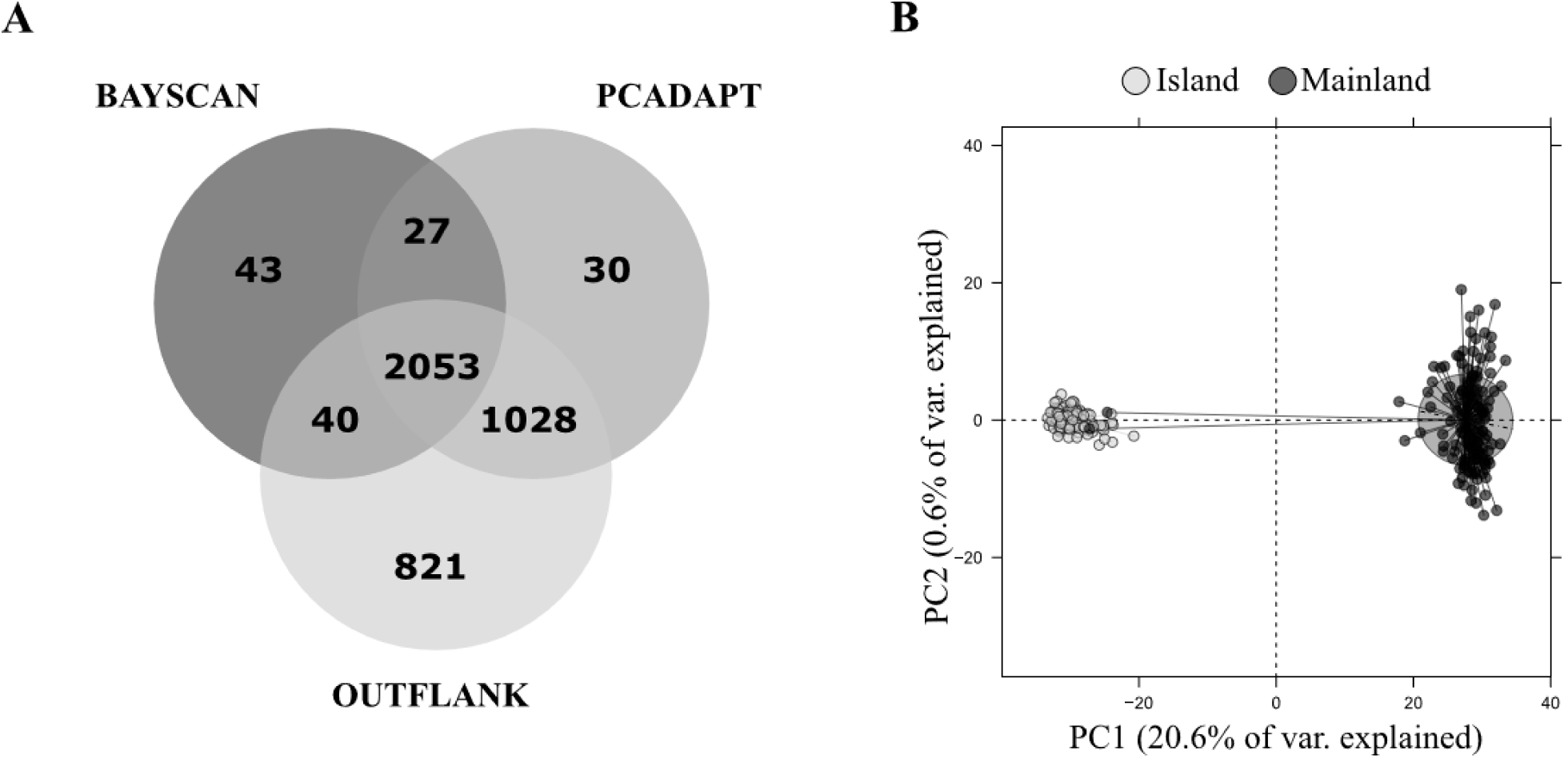
(A)Venn diagram showing the number of putatively unique and shared adaptive SNPs detected by BAYESCAN, PCADAPT, and OUTFLANK. (B) Principal component analysi based on all 2,053 shared candidate SNPs for Torrey pine individuals, including individuals from both mainland (black, n = 140) and island (grey, n = 130) populations. Variation explained by the first two principal components is provided in parentheses.

Functional categorization of common outlier loci was performed by blasting *de novo* assembled contigs carrying outlier SNPs against the *Pinaceae* UniProt protein database and retrieving each hit’s Gene Ontology terms. Overall, 110 (7.51%) contigs were annotated. After accounting for redundancy in the data (i.e., different contigs aligning to the same locus), we identified a total of 80 putative adaptive genes with homologous sequences in *Larix*, *Picea*, or *Pinus* species (Appendix S8) that may be targets of selection. Functionally, these genes are primarily encoded in mitochondria, involved in the process of DNA integration, or with processes associated with molecular functions such as RNA-directed DNA polymerase activity or nucleic acid binding (Table 3).

**Table 3.** Functional categorization of 80 putatively adaptive genes between Torrey pine populations. Listed are the ten most frequent GO terms within each of the three GO classes (Biological Process, Molecular Function, Cellular Component).

| GO class | GO term | Frequency (%) |
| --- | --- | --- |
| Biological Process | DNA integration | 6.2 |
|  | Methylation | 2.5 |
|  | Actin filament polymerization | 1.2 |
|  | Arp2/3 complex-mediated actin nucleation | 1.2 |
|  | Carbohydrate metabolic process | 1.2 |
|  | Carbohydrate transport | 1.2 |
|  | Cell differentiation | 1.2 |
|  | Defense response to bacterium | 1.2 |
|  | Defense response to fungus | 1.2 |
|  | Gene silencing by RNA | 1.2 |
| Molecular Function | RNA-directed DNA polymerase activity | 11.2 |
|  | Nucleic acid binding | 7.5 |
|  | ATP binding | 3.8 |
|  | ADP binding | 2.5 |
|  | Magnesium ion binding | 2.5 |
|  | O-methyltransferase activity | 2.5 |
|  | Oxidoreductase activity | 2.5 |
|  | Protein kinase activity | 2.5 |
|  | 4-lactone oxidase activity | 1.2 |
|  | Actin binding | 1.2 |
| Cellular Component | Mitochondrion | 38.8 |
|  | Integral component of membrane | 7.5 |
|  | Arp2/3 protein complex | 1.2 |
|  | Cell wall | 1.2 |
|  | Chloroplast | 1.2 |
|  | Chloroplast thylakoid membrane | 1.2 |
|  | Cytoplasm | 1.2 |
|  | Extracellular region | 1.2 |
|  | Membrane | 1.2 |
|  | Nucleus | 1.2 |

## Discussion

An understanding of demographic and adaptive evolutionary history is invaluable for rare species of conservation concern, particularly when management decisions impact populations at risk of extinction. Teasing apart the contribution of both stochastic and deterministic evolutionary processes to population genomic differentiation over time and space can be used to inform species conservation decisions, including the potential consequences of genetic rescue (Frankham et al., 2011; Hufford & Mazer, 2003; Ralls et al., 2018). Here, we evaluated the genomics of Torrey pine, a critically endangered endemic isolated to two populations in California. We modeled demographic change and connectivity over time and tested the influence of neutral and selective processes have had on contemporary population genomic structure. We observed that Torrey pine populations exhibited exceedingly low genetic variation, particularly for a conifer (see below), with little within- or among-population structure. Although some connectivity has been maintained between island and mainland populations, demographic modeling indicates that Torrey pine has consistently suffered from low effective population size. Genome scans revealed over 2000 loci that were candidates for selection. Consistent with previous observations indicating phenotypic differences between island and mainland populations, these data suggest adaptive genetic differences have evolved among populations (Di Santo et al., *in review*; Haller, 1986; Hamilton et al., 2017). From a conservation standpoint these results lead to contrasting recommendations with respect to genetic rescue. A history of reduced effective population size and low genome-wide differentiation at neutral loci indicate little genetic differentiation among populations that may impact a genetic rescue program. However, previous observations of phenotypic differences paired with loci associated with divergent selection point towards the importance of adaptive evolution among Torrey pine populations. These results suggest increased genetic variation via inter-population crosses may be needed in this species, but admixture should be evaluated first to quantify its fitness impact.

### Standing genetic variation

Torrey pine populations exhibited extremely low genetic variation in comparison with other conifers which often exhibit an expected heterozygosity around 0.3 within populations (e.g., Namroud et al., 2008; Tsumura, Uchiyama, Moriguchi, Ueno, & Ihara-Ujino, 2012). Average expected heterozygosity (H_E_) and contemporary effective population size (Ne) were estimated at 0.185 and 2,305 individuals for the island population, and 0.184 and 1,715 individuals for the mainland population, respectively (Table 2; Fig. 5). Although low, this study is the first to find genetic variability within island and mainland populations of Torrey pine. While previous genetic analyses using allozymes and chloroplast DNA markers identified fixed genetic differences between populations, they failed to observe genetic variation within populations (Ledig & Conkle, 1983; Whittall et al., 2010). Ledig & Conkle, (1983) hypothesized reduced genetic diversity was attributable to drastically reduced mainland and island populations in the recent geological past. They suggested that Torrey pine populations declined to fewer than 50 individuals, following which a recovery led to approximately 3,000 to 4,000 reproductively mature trees (J. Franklin & Santos, 2011; Hall & Brinkman, 2015). Demographic models herein, however, suggest that effective population size has always been low for Torrey pine (Fig. 5). The best fit demographic model predicted an ancestral effective population size (N_A_) of 1,124 and only limited change following population divergence (Fig. 5). Given these observations, long-term reduced effective population size has likely exacerbated the consequences of genetic drift leading to an extreme lack of genetic variation within Torrey pine populations.

Despite extremely low genetic variation and small effective population sizes, there was no evidence for inbreeding (F_IS_) or excessive relatedness (θ) within populations (Table 2). These findings support high ratios between effective and census population size (Ne/N) found in both Torrey pine populations (island = 0.75, mainland = 0.45) and may, at least partly, explain why these estimates were higher than ratio averages of 0.1 to 0.2 typically recommended for conservation management (Frankham et al., 2014). Indeed, it is not uncommon for plants to exhibit Ne/N ratios greater than 0.2 (Hoban et al., 2020; Waples, Luikart, Faulkner, & Tallmon, 2013). Combined with a lack of within population genetic structure (Fig. 4, Appendix S7), these results also indicate that neither reproduction among relatives nor unequal reproductive success has likely contributed to reduced genetic variation within populations. Wind pollination and zoochorous seed dispersal have likely contributed to homogenizing the gene pool within populations (M. Johnson, Vander Wall, & Borchert, 2003; Loveless & Hamrick, 1984). Pines also possess mechanisms that can reduce the probability of self-fertilization, including the embryo lethal system (Williams, 2009; Williams, Zhou, & Hall, 2001). This self-incompatibility system selectively induces death in embryos resulting from self-fertilization (Bramlett & Popham, 1971; Williams, Barnes, & Nyoka, 1999). Consequently, increased dispersal potential paired with post-zygotic barriers limiting the probability of mating among relatives have likely reduced within-population genetic structure in Torrey pine.

### Neutral genetic differences across populations over time

Demographic modelling using neutral genomic variation supports the maintenance of some genetic connectivity following population divergence approximately 1.2 MYA (Fig. 5), estimating the probability of gene exchange at 8.34×10^−3^ per generation. Despite geographic isolation among populations and reduced potential for inter-population gene flow, contemporary estimates of F_ST_=0.013 indicate only subtle genome-wide differentiation between island and mainland populations. Reduced genetic differentiation is typical of many conifers (Eckert et al., 2010; Namroud et al., 2008; Tyrmi et al., 2020), as pollen may maintain connectivity over very long distances (Campbell, McDonald, Flannigan, & Kringayark, 1999; Varis, Pakkanen, Galofré, & Pulkkinen, 2009; Williams, 2010). Gene flow between populations may also have been maintained via seed dispersal. Birds represent potential seed dispersers for Torrey pine and may play a prominent role in long-distance seed dispersal (M. Johnson et al., 2003; Pesendorfer, Sillett, Koenig, & Morrison, 2016; Viana, Gangoso, Bouten, & Figuerola, 2016). Interestingly, higher estimates of contemporary effective population size (N_I_, N_M_) relative to the ancestral population size (N_A_) suggest that island and mainland populations have experienced genetic bottlenecks following one or multiple moderate population expansion events (Fig. 5). Overall, our findings indicate that despite the increased probability of genetic drift due to genetic bottlenecks and low population sizes, gene flow maintained between island and mainland populations may have been sufficient to prevent extensive genomic differentiation at neutral loci following population isolation. Note, however, that coalescent simulations assume non-overlapping generations, which may limit their ability to accurately estimate demographic parameters in long-lived species, including conifers. Consequently, gene flow estimated between island and mainland populations may have been overestimated or may possibly be an artefact resulting from an attempt of the demographic model to account for shared ancestral genetic variation among populations.

### Evidence for local adaptation

Phenotypic monitoring using common garden experiments or *in situ* morphological observations for cone, seed, and needle morphology have previously suggested genetically-based phenotypic divergence among Torrey pine populations (Di Santo et al., *in review*; Haller, 1986; Hamilton et al., 2017). Thus, subtle genetic differentiation observed among populations likely reflects the fact that most genome divergence among populations is the result of neutral rather than adaptive differentiation among populations. To test for the role of selection across loci, we simulated a null F_ST_ distribution to compare with our empirical F_ST_ distribution, which indicated that a few thousand loci may be under divergent selection (Appendix S5). Genome scans further supported this observation, identifying 2,053 (2.21%) candidate SNPs with accentuated divergence between island and mainland populations (Fig. 6). Annotation of sequences containing these outliers SNPs suggested that adaptive evolution in Torrey pine may not only result from genetic differentiation at the nuclear level, but also at the mitochondrial level (Table 3). This could be consistent with previous observations of the importance of cytoplasmic genetic differences as a factor contributing to local adaptation in plants (Hamilton & Aitken, 2013; Leinonen, Remington, Leppälä, & Savolainen, 2013; Leinonen, Remington, & Savolainen, 2011; for a review see Bock, Andrew, & Rieseberg, 2014). For example, Leinonen et al., (2011) found using a reciprocal transplant experiment that individuals of *Arabidopsis lyrata* harboring the local cytoplasmic genome had higher fitness than individuals harboring the non-local cytoplasmic genome, suggesting that cytoplasmic genetic variation may contribute to local adaptation.

Overall, GO annotation of both nuclear and mitochondrial encoded candidate genes indicated that genes important for mechanisms such as DNA integration, methylation, gene silencing, carbohydrate transport and metabolic processes, and defense against pathogens (bacteria and fungi) were candidates for selection (Table 3). This suggests that between the island and mainland environments, modification of genetic composition and architecture following DNA integration, changes in gene expression or protein function following methylation and gene silencing, and direct or indirect selection against pathogens may have played an important role in population divergence following isolation. For example, a candidate gene associated with defense against bacteria and fungi (UniProt accession: B8LLJ5, GO terms: GO:0042742, GO:0050832, GO:0031640) suggests that phenotypic differentiation may have evolved in response to pests or pathogens. Indeed, the mainland population of Torrey pine may have faced substantial selection associated with the recent outbreak of the California five-spined ips (*Ips paraconfusus* Lanier) (J. Franklin & Santos, 2011; Shea & Neustein, 1995), whereas the island population may not have been exposed to that selective pressure. Noteworthy with these results is the fact that pine genomes are enormous (Grotkopp, Rejmánek, Sanderson, & Rost, 2004; Stevens et al., 2016), and our sequencing approach (reduced representation sequencing; see Material and Methods) represents only a fraction of the Torrey pine genome. This suggests, despite the differences observed, most likely some variation has been overlooked that plays a critical role in local adaptation for this species.

### Applying neutral and adaptive evolutionary processes to rare species conservation

While identification of the appropriate effective population size necessary to protect adaptive evolutionary potential for rare species is still debated, recommendations generally range between 500 to 5000 individuals (Frankham et al., 2014; I. R. Franklin & Frankham, 1998; Lynch & Lande, 1998). Torrey pine, critically endangered and endemic to just two native populations, suffers from extremely low effective population size (N_I_ = 2,305, N_M_ = 1,715) relative to other pines (Menon et al., 2018; Xia et al., 2018). In addition, there is clear evidence of the impact of adaptive evolutionary processes alongside neutral processes structuring genomic variation within and among populations. Given historical and contemporary estimates of effective population size as well as contemporary estimates of expected heterozygosity, our results indicate that Torrey pine may not retain the genetic variation within populations needed to adapt to change. Current monitoring within the mainland population suggests that a lack of recruitment (personal observation), infestation by *Ips* beetles (personal observation), and climate warming (Diffenbaugh, Swain, & Touma, 2015) may increase extinction risk. Thus, for Torrey pine, increased Ne and greater genetic diversity may be required for long-term persistence.

As genetic variation is extremely low within populations, one conservation strategy that may facilitate the maintenance of genetic variation within populations at risk is a genetic rescue program. A genetic rescue program would facilitate inter-population breeding as a means to increase heterozygosity, increasing rates of inter-population gene flow. Indeed, demographic modeling indicates that following population isolation some gene flow has been maintained and there are low levels of genome-wide differentiation among populations (Nei’s F_ST_ = 0.013). However, the combination of observed phenotypic differences and large number of genes that appear targets of selection suggest island and mainland populations of Torrey pine have undergone distinct evolutionary trajectories necessary for adaptation following isolation. Thus, a genetic rescue program should be considered with caution as gene flow between populations may disrupt local adaptation and further reduce population performance (Goto, Iijima, Ogawa, & Ohya, 2011; Hufford & Mazer, 2003; Montalvo & Ellstrand, 2001). Despite this word of caution, preliminary data comparing mainland, island, and F1 individuals from a common garden experiment planted outside the species natural distribution indicate that F1s exhibit increased fitness relative to mainland and island populations (Hamilton et al., 2017). Consequently, future monitoring is needed to empirically quantify fitness consequences of advanced-generation admixture (F2, Backcross-Island (BC-I), Backcross-Mainland (BC-M)) following early-generation heterosis.

Given the challenge to conserve and manage rare species in a rapidly changing environment, the use of genomic data to model evolutionary history, assess demographic change, and tease apart the contributions of neutral and adaptive processes will be critical. For Torrey pine, the fact that there is low genome-wide differentiation among populations, a consistent history of low effective population size, and indications that some gene flow is maintained among populations may suggest that one population (island or mainland) could be targeted to effectively preserve neutral genetic variation. However, the combination of outlier loci and previously observed phenotypic differences suggest if the goal is to preserve adaptive genetic variation, a strategy that favors conservation efforts across both mainland and island populations will be needed. If conservation strategies such as genetic rescue are considered, assessment of multiple admixed generations within a common environment will provide the necessary empirical test to evaluate the consequences of enhancing genetic exchange among populations.

## Acknowledgements

This work was funded by USDA Health Protection Gene Conservation program and Western Wildlands Environmental Threat Assessment Center to JAH and JWW, Morton Arboretum Center for Tree Science Fellowship to JAH, and a new faculty award from the office of North Dakota Experimental Program to Stimulate Competitive Research (ND-EPSCoR NSF-IIA-1355466) and the NDSU Environmental and Conservation Sciences Graduate Program to JAH and LNDS. For their help collecting needle tissue, the authors would like to thank Stephen Johnson, Jill Wulf, Forest Swaciak, Conner Harrington, Emma Ordemann, Drew Peterson, Andrew Bower, Gary Man, Valerie Gallup, Annette Delfino Mix, Stephanie Steel, and tree climbers from San Diego Zoo Wildlife Alliance. For their help coordinating sampling on Santa Rosa Island and at the Torrey Pine State Reserve, the authors also would like to thank Paula Power (NPS), Rocky Rudolph (NPS), Kathryn McEachern (USGS), Robyn Shea (California State University – Channel Islands), and Darren Smith (California State Parks). Finally, the authors would like to acknowledge the Michumash and the Kumeyaay people as the traditional caretakers of the Torrey pine ecosystems sampled for this study. Any use of product names is for informational purposes only and does not imply endorsement by the US Government.

