## Supplemental Material S1-S8 for "Reduced representation sequencing to understand the evolutionary history of Torrey pine (*Pinus torreyana* Parry) with implications for rare species conservation"

and Jill A. Hamilton

**Appendix S1.** Distribution of genetic summary statistics in the HWE-filtered (73,928 variants, dark grey) and down-sampled (9,795 variants, light grey) datasets. (A, C) observed heterozygosity: H_O_. (B, D) pairwise genetic distances: Nei’s F_ST_.

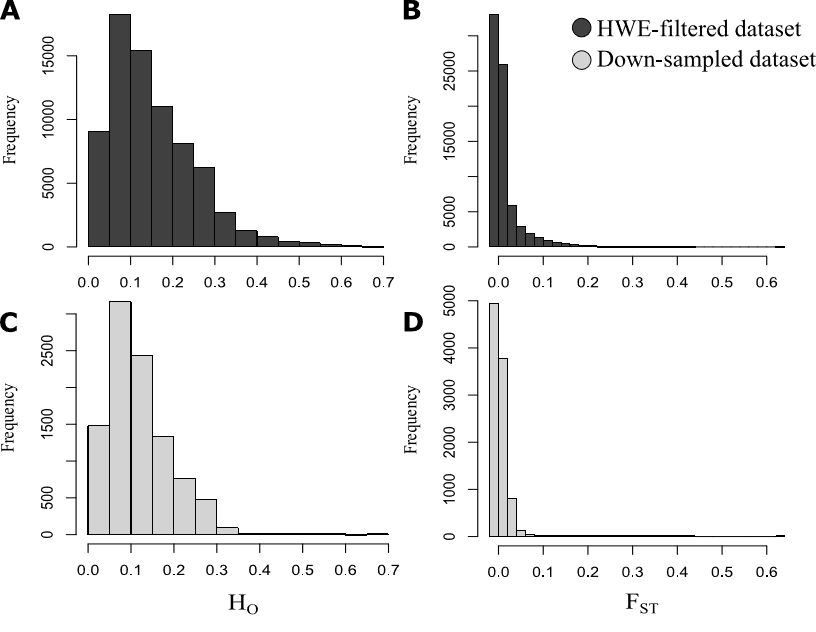

**Appendix S2.** Principal component analyses conducted on two different genetic datasets. (A) HWE-filtered dataset, including 73,928 SNPs. (B) Down-sampled dataset, including 9,795 SNPs. Individuals were grouped by populations, including island (SRI, grey) and mainland (TPSR, black) populations. Variance explained by the first and second axis of differentiation are provided in parentheses.

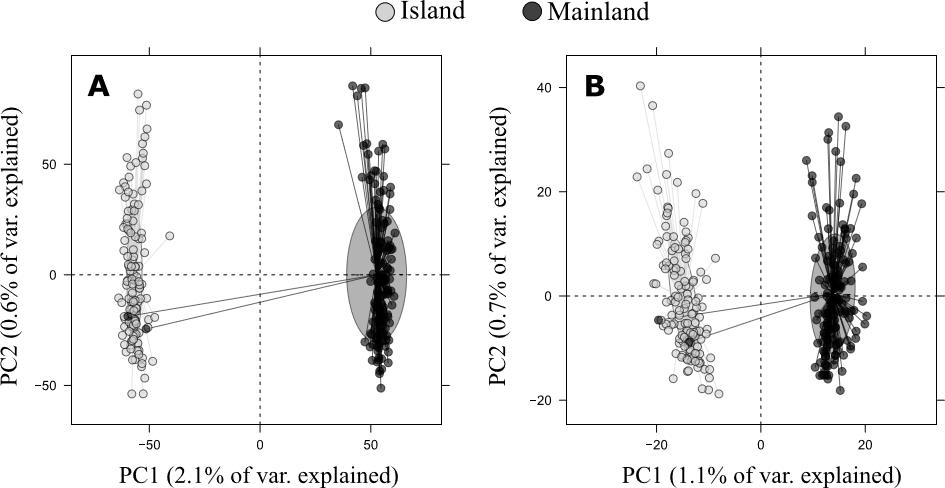

**Appendix S3.** Misclassification proportions based on 100-fold cross-validation simulations with a tolerance rate of 0.01 for all six demographic scenarios. Different colors indicate summary statistics simulated under different demographic scenario. The proportion of simulations correctly assigned to their demographic model is provided above each bar. See text and Figure 2 for details on each scenario.

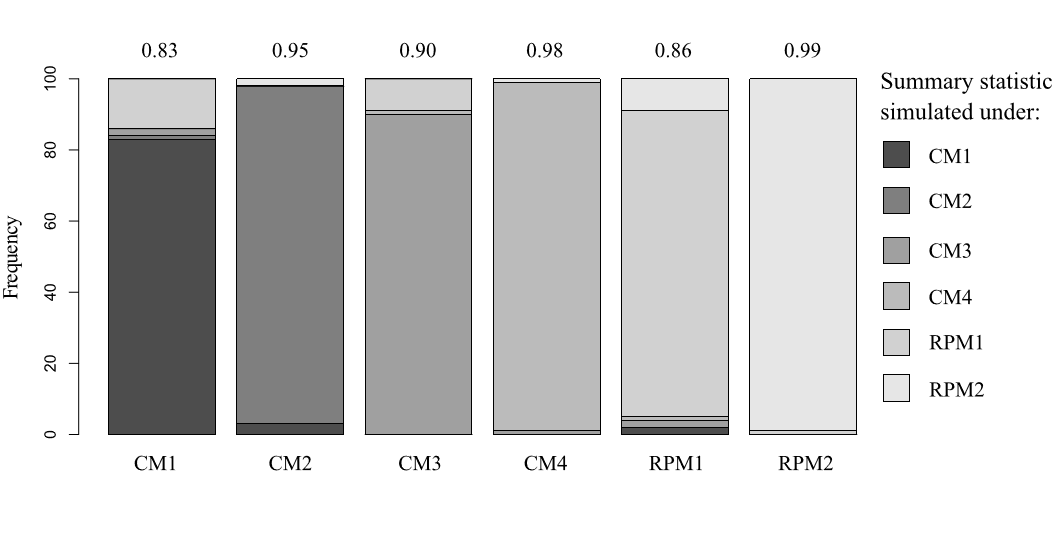

**Appendix S4.** Cross-validation (100-fold) for demographic parameters inferred from the best model. True values (x-axis) are given against estimated values (y-axis) approximated using non-linear postsampling regression adjustment on log-transformed parameters and a tolerance rate of 0.01. The solid line represents the 1:1 (identity) relationship between true and estimated values. The closer to the line points are, the more accurate is the parameter inferred. N_A_, ancestral effective population size; N_I_, island effective population size; N_M_, mainland effective population size; m_IM_, migration probability from island to mainland; m_MI_, migration probability from mainland to island; T_Div_, time of population divergence.

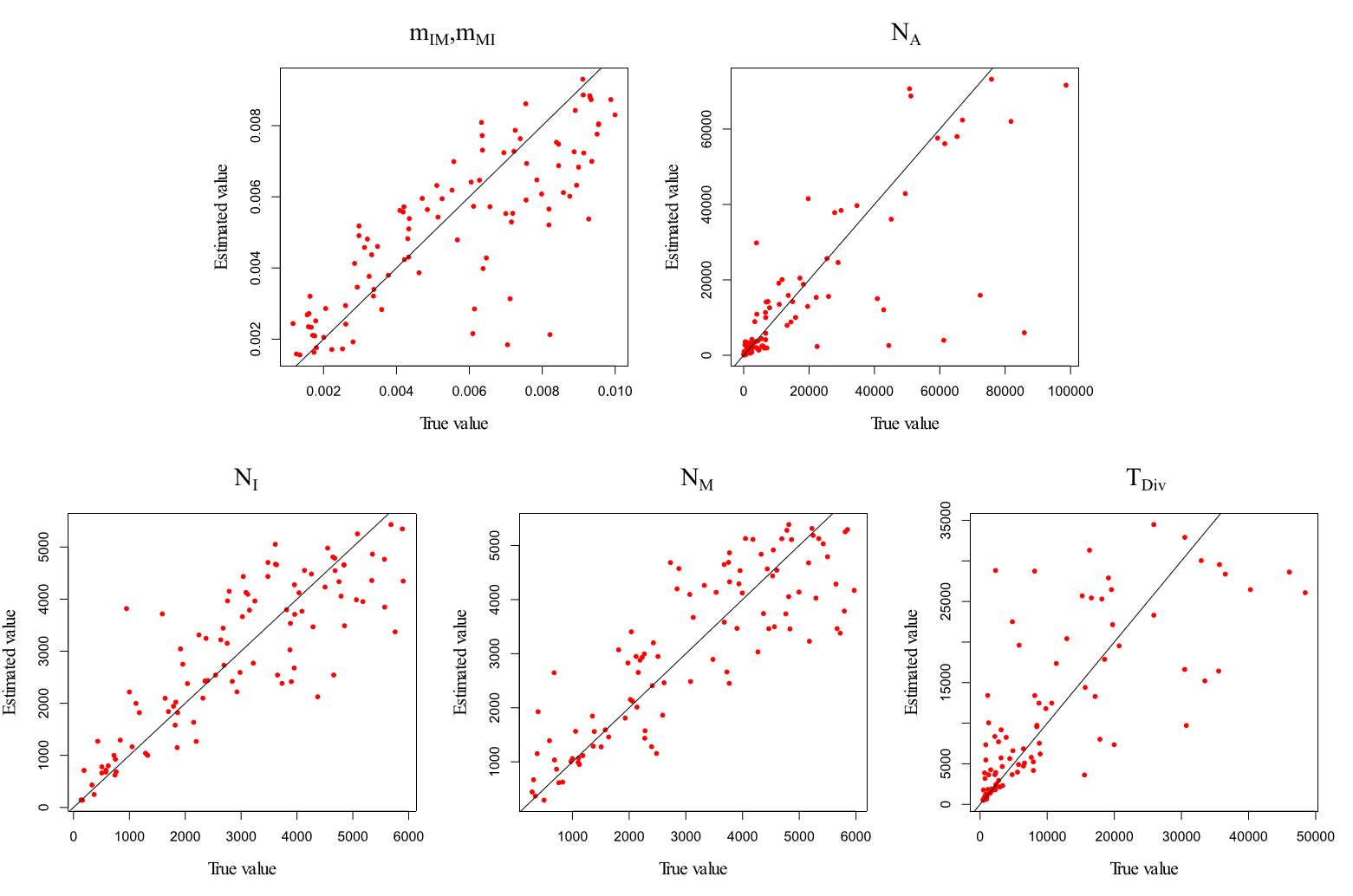

**Appendix S5.** Distributions of pairwise Nei’s F_ST_ values between the island and mainland populations estimated from the simulated (grey) and full (white) dataset. Both distributions were estimated using 93,085 SNPs. (A) Full range of observed and simulated F_ST_ values. (B) Right-hand tail of observed and simulated F_ST_ distributions. Neutral F_ST_ estimates were obtained by simulating the best demographic model (see Figure 5) in SIMCOAL2.

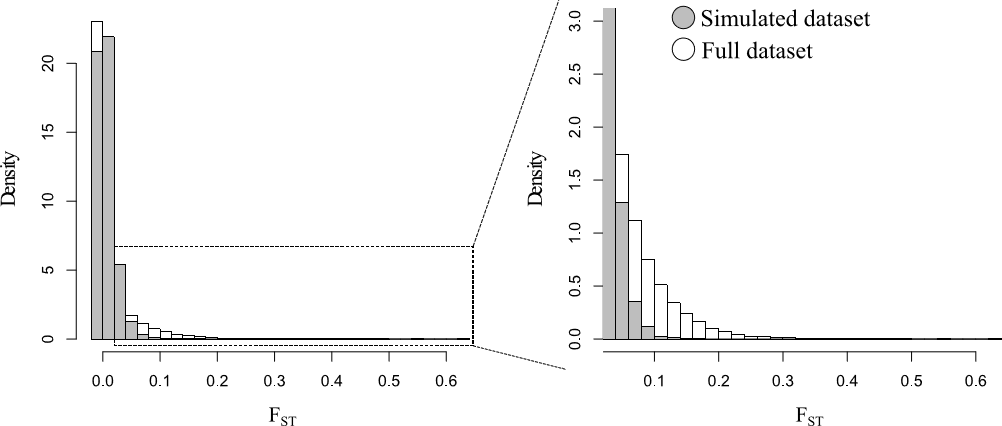

**Appendix S6.** Principal component analysis conducted on the full dataset (93,085 variants) grouped by populations (island [SRI], grey; mainland [TPSR], black). (A) Distribution of individuals on the first and second axis of differentiation. (B) Distribution of individuals on the third and fourth axis of differentiation. Variation explained by each axis is given in parentheses.

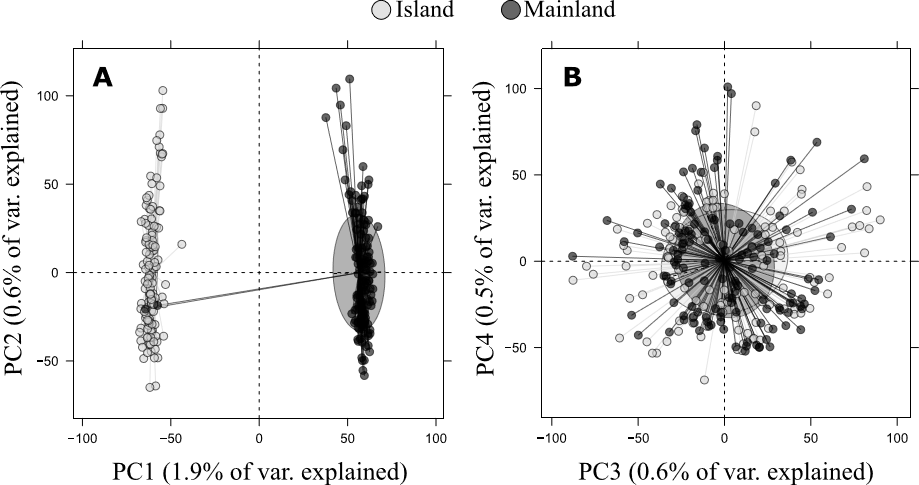

**Appendix S7.** Bayesian Information Criterion (BIC) calculated for structure models assuming from 1 to 10 genetic clusters (*k*) within Torrey pine populations. (A) Island (SRI) population. (B) Mainland (TPSR) population. For both populations, *k* = 1 is associated with the lowest BIC value, identifying one as the most likely number of genetic clusters within populations.

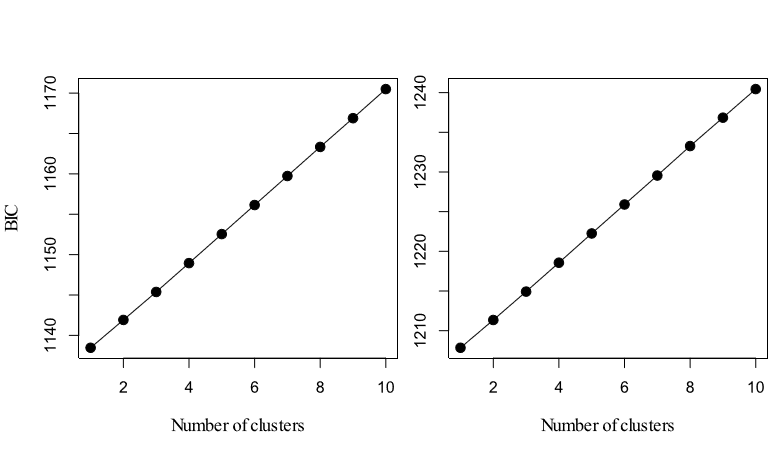

**Appendix S8.** *Pinaceae* species represented in the set of 80 putatively adaptive genes.

| **Species**  **with BLAST’s hits** | **Number of**  **BLAST’s hits** |
| --- | --- |
| *Larix gmelinii* | 3 |
| *Larix kaempferi* | 1 |
| *Picea abies* | 3 |
| *Picea glauca* | 26 |
| *Picea sitchensis* | 25 |
| *Pinus elliottii* | 1 |
| *Pinus lawsonii* | 1 |
| *Pinus massoniana* | 4 |
| *Pinus monticola* | 1 |
| *Pinus pinaster* | 2 |
| *Pinus pinea* | 1 |
| *Pinus radiata* | 3 |
| *Pinus sylvestris* | 2 |
| *Pinus tabuliformis* | 2 |
| *Pinus taeda* | 5 |
| **Total** | **80** |
